# Viroid-sized rings self-assemble from mononucleotides through wet-dry cycling: implications for the origin of life

**DOI:** 10.1101/2020.04.28.064972

**Authors:** Tue Hassenkam, Bruce Damer, Gabriel Mednick, David Deamer

**Affiliations:** Nano-Science Center, Department of Chemistry, University of Copenhagen, DK-2100, Denmark; Department of Biomolecular Engineering, University of California, Santa Cruz CA 95064, California, USA; UpRNA, Santa Cruz CA 95060, USA; Department of Chemistry and Biochemistry, University of California, Santa Cruz CA 95064, California, USA

## Abstract

It is possible that early life relied on RNA polymers that served as ribozyme-like catalysts and store genetic information^1,2^. The source of such polymers is uncertain, but previous investigations reported that wet-dry cycles simulating prebiotic hot springs provide sufficient energy to drive condensation reactions of mononucleotides to form oligomers^3^. The aim of the study reported here was to visualize the products by atomic force microscopy. In addition to globular oligomers, ring-like structures ranging from 10-200 nm in diameter with an average around 30-40 nm were abundant, particularly when nucleotides capable of base pairing were present. The thickness of the rings was consistent with single stranded products, but some had thicknesses indicating base pair stacking. Others had more complex structures in the form of short polymer attachments and pairing of rings. These observations suggest the possibility that template-directed synthesis may occur during wet-dry cycling followed by solvation of the rings. We conclude that RNA-like rings and structures could have been synthesized non-enzymatically on the prebiotic Earth with sizes sufficient to fold into ribozymes and genetic molecules required for life to begin.

---

Our current understanding of the early Earth suggests that life originated approximately four billion years ago^4-6^, and that early stages of life depended on polymers resembling RNA^1,2^. Organic compounds from geochemical sources and meteoritic infall^7^ likely accumulated on volcanic land masses emerging from a global ocean. Some were flushed by precipitation into hydrothermal pools supplied with fresh water distilled from a salty ocean^3,8^. Water levels in some pools fluctuated due to variation in the flow of hot spring water, geyser activity and precipitation. Solutes would be concentrated by evaporation and form films on hot mineral surfaces in which condensation reactions synthesize biologically relevant polymers^3,8,9^. For instance it has been shown that short oligomers linked by peptide bonds are synthesized from mixtures of amino acids during wet-dry cycling^10-12^. More recently it has been reported that ribonucleotides can be synthesized from simple organic compounds in conditions that incorporate a wet-dry cycle^13,14^.

If so, nucleotide monomers present as solutes in the pools may also have formed polymers under these conditions on the prebiotic Earth. In previous studies, we have reported a series of experimental results supporting the conclusion that such conditions can drive polymerization of mononucleotides:

1. Oligomers produced by wet-dry cycling of mononucleotides form pellets when isolated by precipitation in 70% ethanol or with spin tubes designed to purify nucleic acids. The oligomers exhibit UV spectra that are characteristic of the mononucleotides composing them.
2. The RNA-like oligomers synthesized by wet-dry cycles are recognized by the enzymes used to label RNA with 32-P. When the labeled material is analyzed by standard methods of gel electrophoresis, it moves through the gel as expected for polyanions and shows up as a band ranging from 20 to >100 nucleotides in length^15^. The oligomers can also bind dyes such as ethidium bromide and SYBR-SAFE that are used to stain RNA polymers in gels^16^.
3. When tested by nanopore analysis with the alpha-hemolysin pore, the oligomers produce blockade signals virtually identical to those caused by single stranded RNA molecules. This demonstrates that at least some of the products are linear polyanionic strands that impede ionic currents as they are driven through the nanopore by an applied voltage^15-17^.
4. If a 1:1 mole ratio of AMP and UMP is exposed to wet-dry cycling, the oligomeric products exhibit hyperchromicity, but the products from AMP alone do not^16^. This result is consistent with hairpin structures forming in random sequence linear polymers of RNA composed of monomers capable of Watson-Crick base pairing.
5. An X-ray diffraction study of AMP-UMP mixtures revealed that linear arrays of stacked bases are present with 3.4 Angstrom distances between the bases^18^. The arrays are referred to as pre-polymers which presumably can form phosphodiester bonds during wet- dry cycles that link them into polymers by condensation reactions.

Although the experimental evidence is consistent with short oligomers being formed during wet-dry cycling^9,15-17,19^, the structure of the oligomers is unknown. We therefore employed atomic force microscopy (AFM) to visualize the products at single molecule resolution.

We exposed dilute solutions of mononucleotides such as adenylic (AMP), uridylic (UMP), guanylic (GMP) and cytidylic (CMP) acid to conditions simulating hot hydrothermal pools undergoing wet-dry cycles. For convenience, hereafter the mononucleotides will sometimes be abbreviated A, U, G and C, and mixtures will be abbreviated AU and GC for 1:1 mole ratios of AMP:UMP or GMP:CMP respectively.

A single cycle is defined here as a 10 mM solution of mononucleotides evaporating on mica (20 - 40 microliters) or glass (100 uL) at 80° C for 30 minutes, then rehydrated by addition of the same volume of water. Because the nucleotides were acids rather than sodium salts, the pH was approximately 2.5. A typical experiment included three such cycles followed by flushing with water to dissolve and remove excess mononucleotides. The mica was then dried for AFM examination.

Figures 1 a and b show a typical result from an AU solution sampled after the last of three wet-dry cycles on a glass substrate. The AFM images are shown in inverted height scale, from light blue to black, the darker the particle the higher it is. We expected to see products appear as small particles of oligomers that had folded into globular forms. Such particles were abundant, but we also observed multiple ring-like structures. The largest was approximately 200 nm in (outer) diameter while the smallest detectable ring was approximately 10 nm in diameter. The average diameter over 100 rings was 39 nm ± 19 nm (more rings are shown in Extended data Fig. 2 and the range is shown in Fig. 3). Smaller rings might have been present in large numbers but due to convolution caused by the tip shape the AFM could not distinguish between a solid particle and rings less than 10 nm in diameter. The rings shown in Figs. 1c and d differ from the rings in Figs. 1 a, b because the wet dry cycle was performed on a freshly cleaved mica surface rather than glass, and the nucleotides were a mixture of GMP and CMP instead of AMP and UMP. The results suggest that rings were produced only when complementary base pairs are present in the solution. The AU and GC rings were similar in thickness, and with similar variations in the integer values of the single layer (cross section in Fig 1c). While most rings were a single layer thick (0.3-0.4 nm) parts of the rings were often multiple layers in thickness suggesting a packing motif with monomers building sequential layers of rings. Multiple examples are found in Fig. 1, We have indicated pairs of rings having the same diameter but with twice the thickness by red arrowheads.

**Figure 1:**
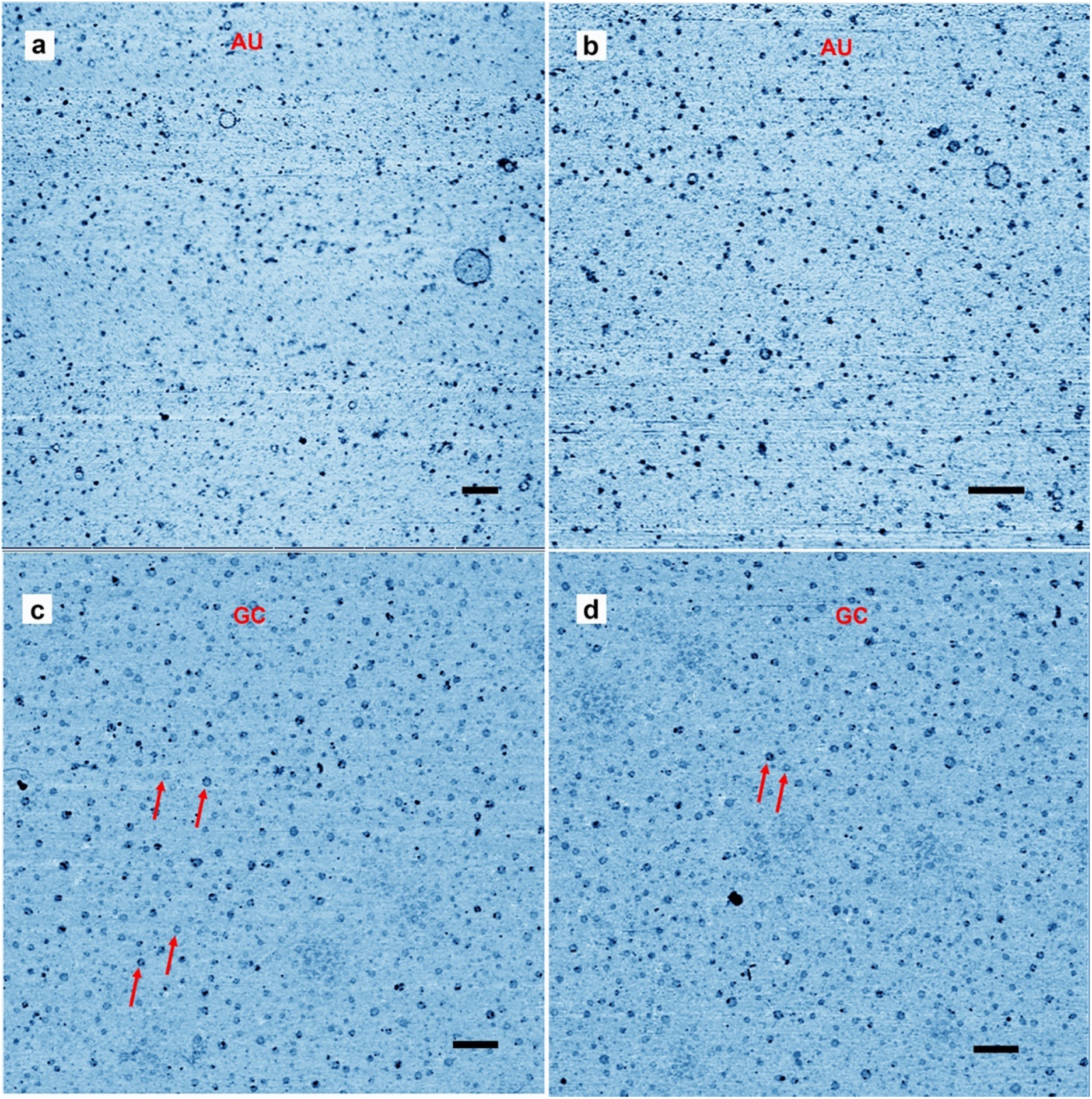
Representative AFM images of rings. A) and B) rings captured on a mica surface from an effluent solution after a hot WD cycle on a glass with AU mixture. The mica surface was exposed to the solution for about 30 seconds as described in the methods section. C) and D) the mica substrate used for hot WD cycles with GC mixture. Scalebars shows 200 nm.

Fig. 2 shows further examples. In Fig. 2a a cross section is drawn across a single layer ring and a ring with partial double layer. In Fig. 2b a cross section is drawn across a full double ring AU and a full double CG ring is shown in Fig 2d. In Fig. 2c the cross section is drawn across a ring with a particle attached to the ring. These types of rings that came decorated with extra material were abundant. There were also many rings that appeared to have small tails of polymeric segments, partial extra layers and small particles attached. Some rings appeared to form pairs either with smaller rings attached to larger rings, or by similar sized rings tethered by short polymer strands. A catalog of these different types rings and structures are shown in Fig. 3.

**Figure 2:.**
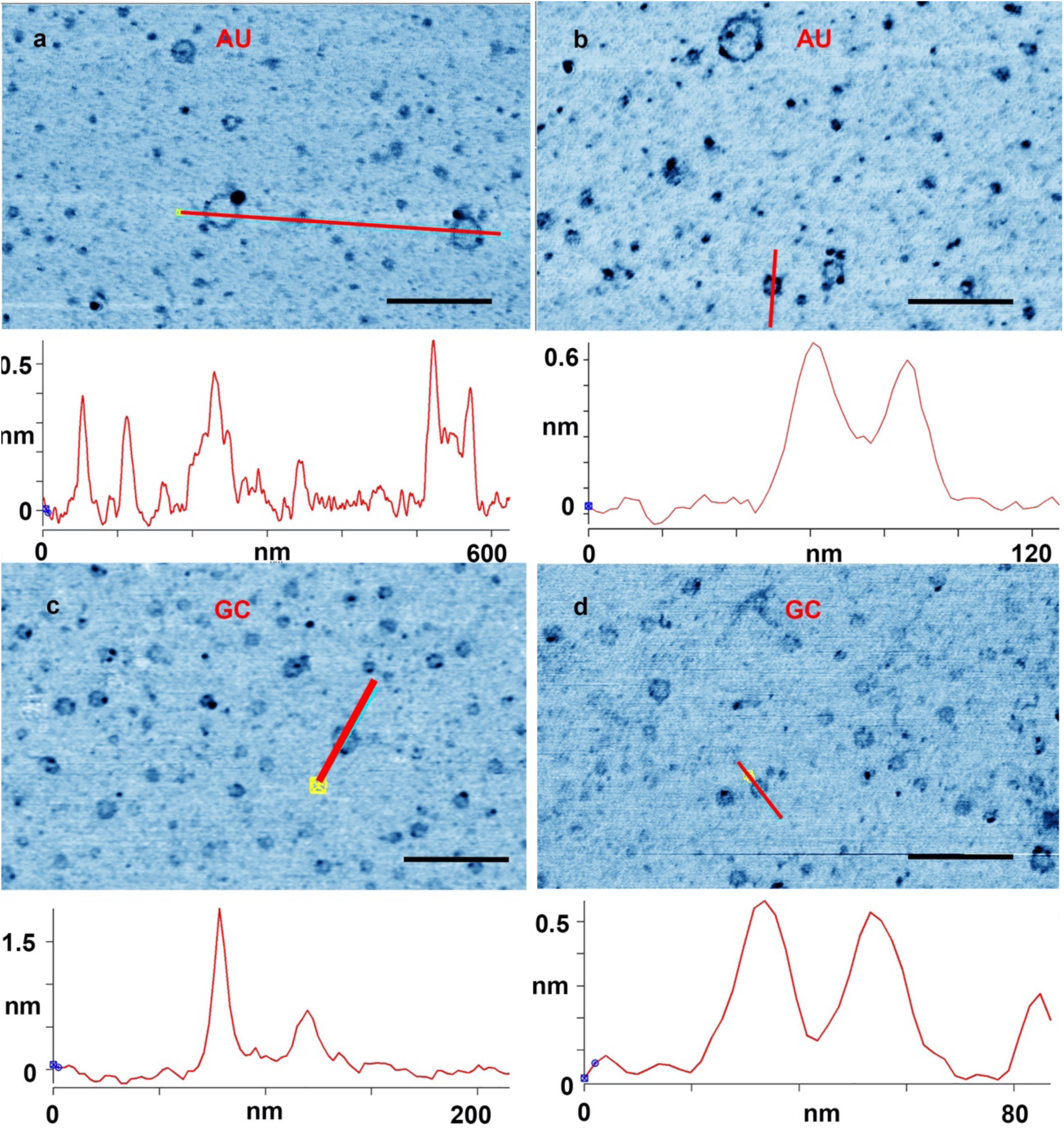
a) and b) rings captured on a mica surface from an AU solution after three wet-dry cycles on glass. c) and d) a mica substrate used for three wet-dry cycles with a GC mixture. The cross sections showing the height profile were drawn along the red lines marked in the images. Scalebars shows 200 nm

**Figure 3:**
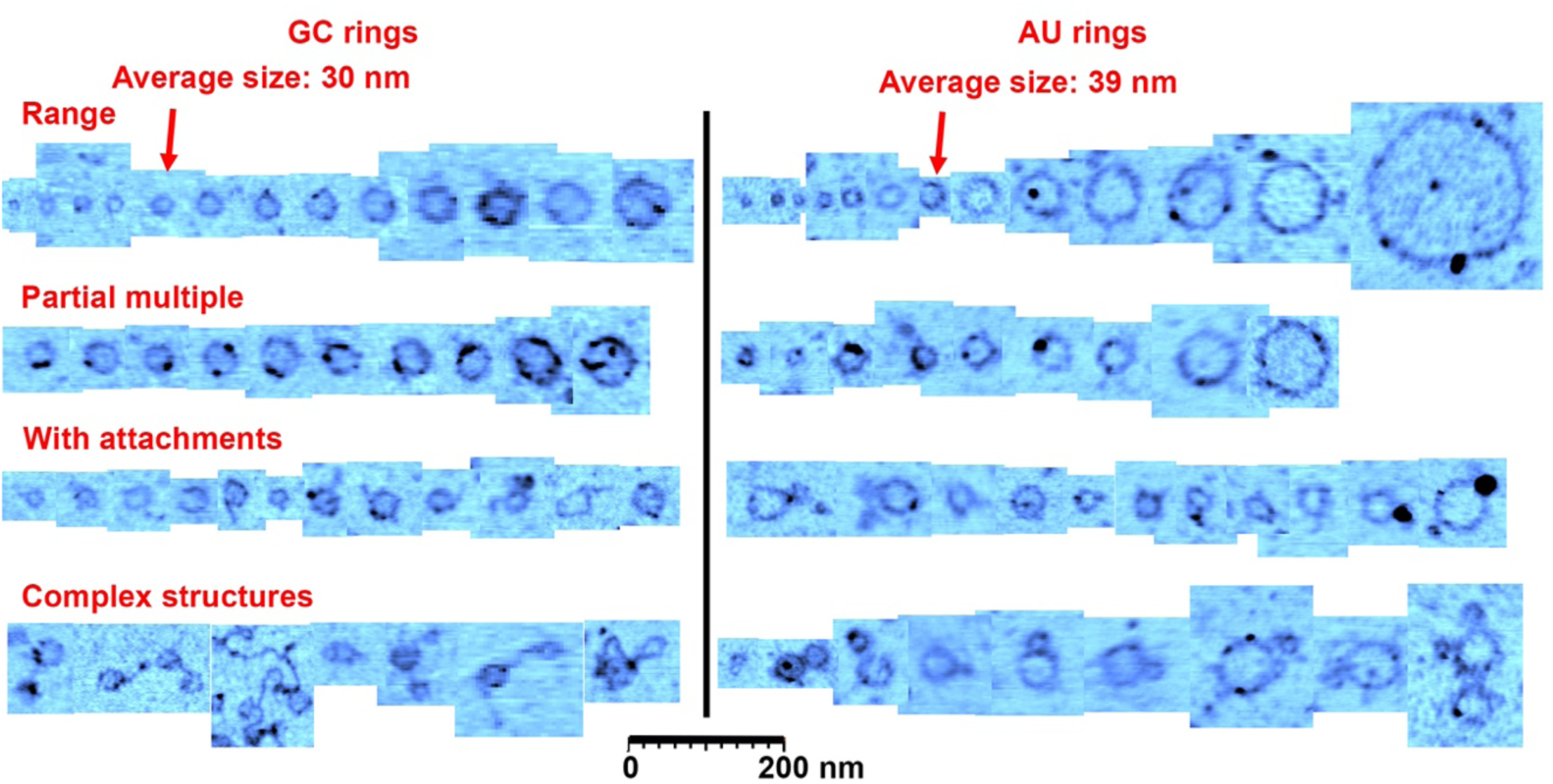
A catalog of ring types. This figure shows the range of ring sizes found on mica samples after wet-dry cycling of GC solutions and AU solutions. Most rings were the size of the average indicated with the arrowheads. The second row shows rings that had segments twice or more in thickness (there is also examples in the other rows), and the third row shows rings with small polymer or particle attachments. The last row shows rare complex structures observed in the images, very often involving several rings attached to each other. The cutouts showing rings and structures were extracted from approximately 30 AFM images.

The observed diameter of the GC rings did not vary as much as the AU rings, with a size distribution ranging from 10 to 60 nm and an average diameter of around 30±7 nm for 1000 rings found in a 12.5 µm^2^ area. (Fig. 1, Extended data Fig. 3).

One explanation for this narrow distribution could be some mechanism in the drying process that dictated the size. In the experiments with nucleoside mixtures that could show if this was the case (Fig. 4), we did observe a few particles in the expected size range, but the number of particles was far less than the number of observed rings in a given area. The rings produced from the AU base pairs displayed a much larger distribution although the average diameter was close to that of the GC rings: 39±19 nm. The AU sample was taken from a solution, however, so it probed a mixture of whatever had dissolved and was adsorbed to the mica surface. Perhaps the size differences were simply due to variations between rings being first desorbed from glass and then adsorbed to mica, and therefore rings from several regions on the glass substrate was sampled, compared to the GC sample where a comparable small region was probed.

**Figure 4:**
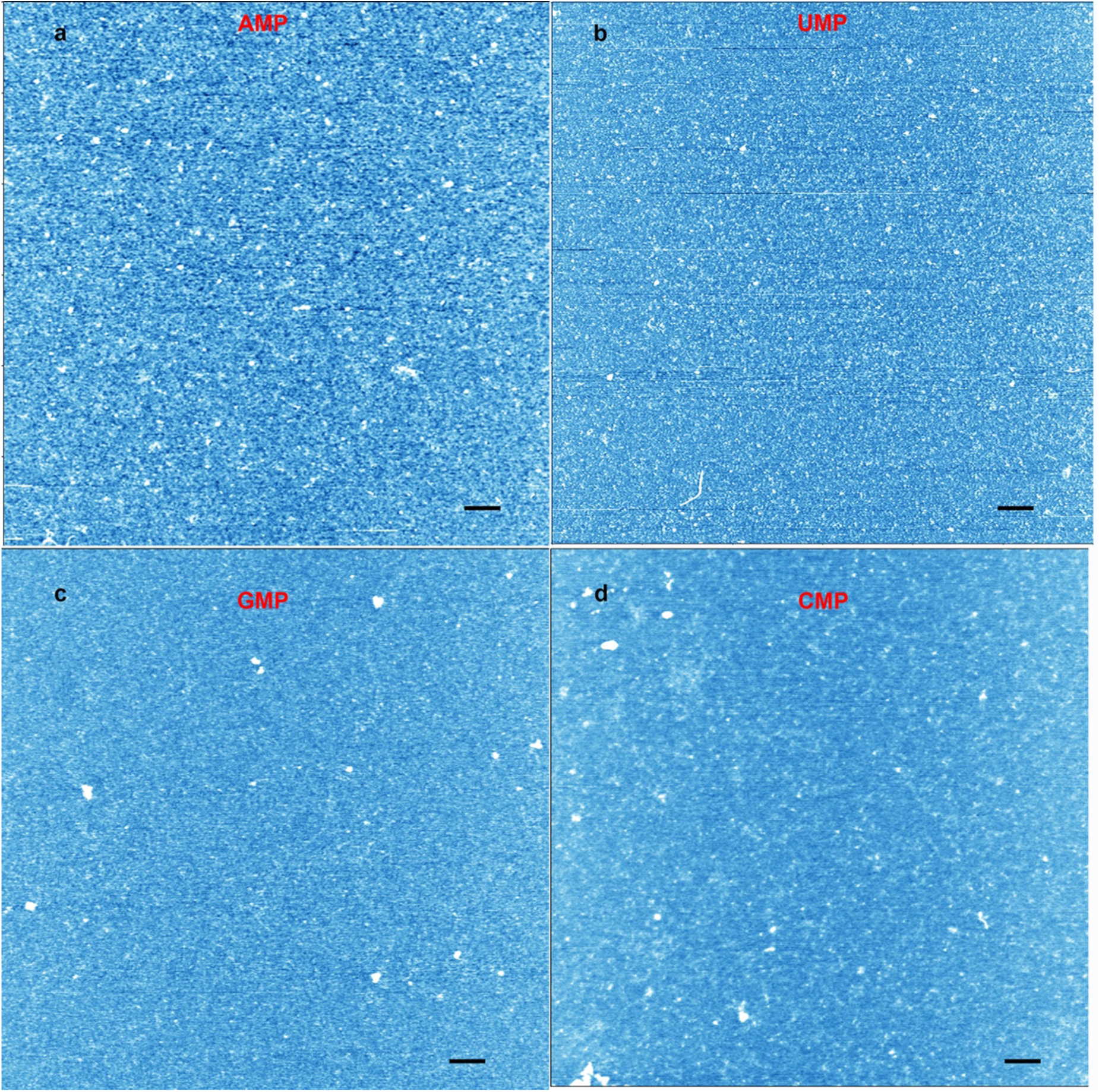
A control experiment was performed with mononucleotide solutions cycled three times on mica. a) AFM image of an AMP solution cycled on mica. b) UMP c) GMP d) CMP. The height scale runs from light blue to white. The scalebars shows 200 nm.

The number of nucleotides present in the rings can be calculated from their diameters (10 - 200 nm) corresponding to 30 - 600 nm in circumference. The number of bases in the rings would then be 50 – 1200 if we use 0.5 nm as the spacing of nucleotides in a single strand, or 70 - 1760 base pairs if the rings are duplex strands having 0.34 nm spacing.

We performed several control experiments to verify that the rings were indeed nucleic acid polymers bound by phosphodiester bonds and not random contamination or artifacts. For instance, we confirmed the presence of rings in two completely independent sets of experiments performed in two different laboratories and repeated the experiments with fresh compounds, solutions, and new tips and surfaces. We tested the mononucleotide solutions to assure that they did not contain contaminating rings before exposure to wet-dry cycles (Extended data 3). We ran wet-dry cycles with individual mononucleotides to see if they also generated rings. Some typical images of mica surfaces cycled with mononucleotides are shown in Fig. 3. Structures consistent with oligomers or short polymers can be observed in all the images, but rings like the ones shown in Fig. 1 and 2 are absent. We also ran wet-dry cycles using two different water sources without mononucleotides and verified that the solvents did not contain compounds that could generate rings.

A second control was to test whether the rings were simply drying artifacts. To this end, we performed wet-dry cycles with matching pairs of nucleosides that lacked phosphate groups. If nucleosides also produced multiple rings, that would argue against the presumed polymerization of nucleotides. We exposed adenosine and uridine, or cytosine and guanosine nucleoside mixtures to wet-dry cycling on mica. We observed a few disk-shaped structures with diameters similar to the rings but these lacked the distinctive ring-shaped perimeters. The disk shape and other structural features are evident from the accompanying AFM amplitude images (Fig. 5 b, d).

**Figure 5:**
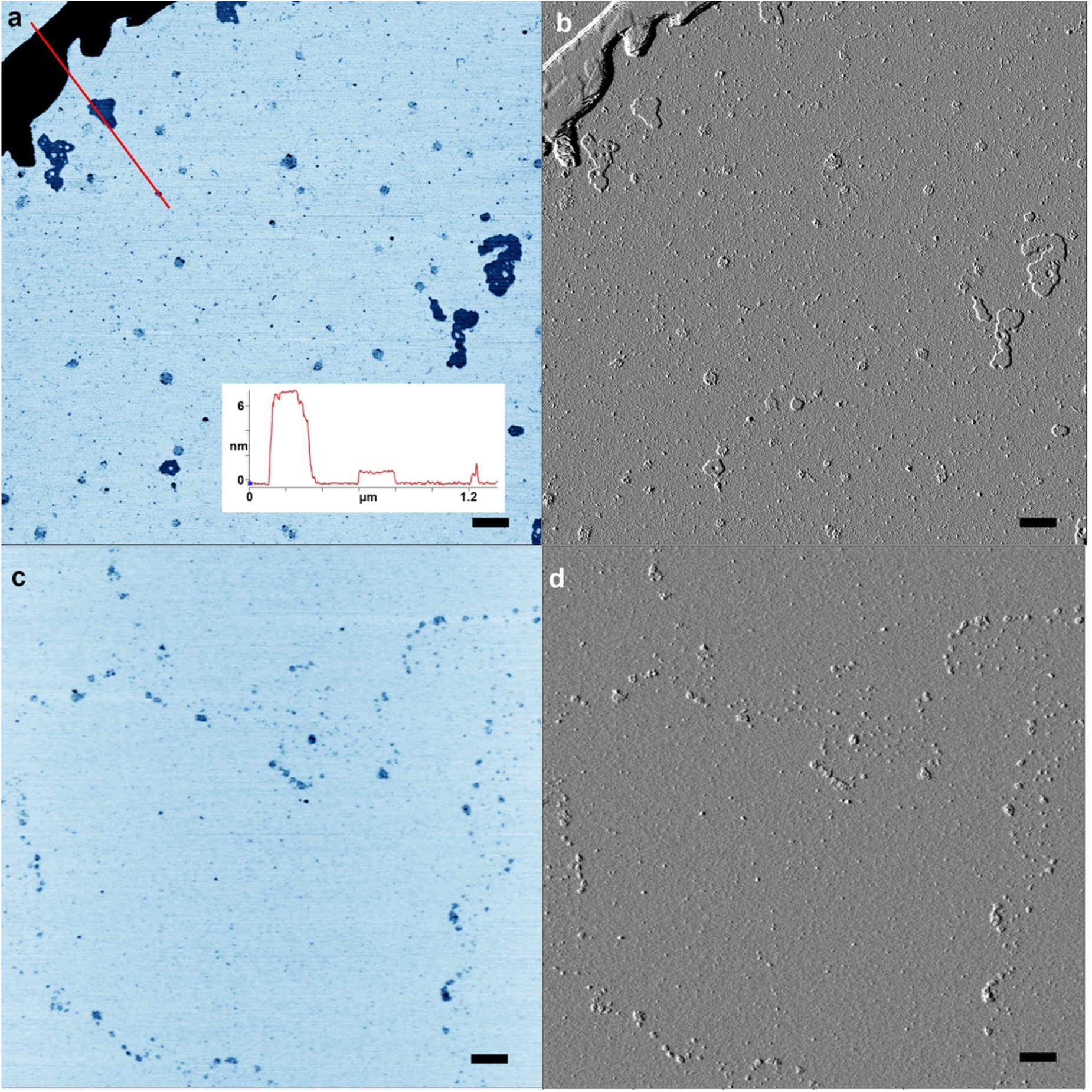
A control experiment was performed with matching pairs of nucleoside mixtures cycled three times on mica. a) AFM image of a 1:1 adenosine:uridine mixture inverted for contrast. b) AFM image of the area shown in a) but using AFM amplitude mode to reveal structures. c) and d) show the same image pairs with a mixture of guanosine and cytidine. The scalebars shows 200 nm

Controls were also run with uncycled 10 mM mononucleotides on mica. The protocol was the same as that used in Fig. 1a, b and Extended data Fig. 1. A few microliters of solution were added to a freshly cleaved mica surface, left there for 30 seconds and then rinsed with pure water. Only particles were apparent, showing that the solutions were not by themselves sources of rings (Extended data Fig. 3).

In another set of controls AU and GC were wet dry cycled at room temperature on a mica substrate. Although none of the controls displayed rings like those observed in Fig. 1, 2, the GC mixture dried at room temperature did self-assemble into rod-like structures (Extended data Fig. 4a) that have been reported previously^20^ It is well known that GMP spontaneously forms tetrameric quadriplexes that assemble into linear rods. Base stacking has also been observed by X-ray diffraction when AU mixtures are dried on various substrates^18^. These are referred to as pre-polymers and may precede polymerization in wet- dry cycling. Rings are created when AU solutions undergo wet-dry cycling on glass surfaces, so mica is not a specific catalytic surface, but we have not excluded the possibility that the slightly negative SiO groups on the silicate surfaces of glass and mica could play a role in the polymerization as has been previously suggested^21,22^

In summary, rings of nucleotides are formed non-enzymatically by ester bond synthesis during wet-dry cycling that simulates conditions present on the prebiotic Earth. The ring structures add to the weight of evidence that the oligomers synthesized by wet- dry cycles resemble RNA in their physical properties^23,24^, and similar ring structures has been observed with DNA^25^. Furthermore, the sizes of the rings are well beyond the minimum range required for catalytic or genetic structures to form. Interestingly they are in the size range of viroid rings composed of 246 - 467 nucleotides, which are the smallest extant infective agents. Viroids were discovered in 1971 by Theodor Diener who speculated later that they may represent RNA remnants of an early form of life^26^. Although the bulk of the GC and AU rings are similar in shape and size, there are some clear variations in terms of size, structural attachments and complexity as shown in Fig. 3. These structural variations could give rise to functioning ribozymes capable of carrying genetic information.

A simple calculation shows that a few micrograms of RNA-like polymers synthesized by wet-dry cycling represent trillions of molecules, each different from all the rest in size and nucleotide sequence. If polymeric rings and their variations of this kind could undergo selection, it would represent the first instance of Darwinian evolution and an early step toward the origin of life as we know it. In a letter to his colleague Joseph Hooker, Charles Darwin speculated that life might begin in a "warm little pond"^27^. The results reported here are consistent with Darwin’s suggestion, with the proviso that the pond must be an acidic solution of monomers undergoing wet-dry cycling at elevated temperatures.

## Acknowledgements

The authors thank Ed Schulak and EdenRoc Sciences, LLC for supporting the study reported here. TH thanks the Villum foundation for support under the “Experiment” program Grant number: 17387, and the Danish Council for Independent Research for support under project 1.

## Authors contributions

TH performed atomic force microscopy, image analysis and the experiments on mica, BD carried out experimental tests within the geological context in which biopolymers assemble from monomers, and GM optimized conditions for nucleotide polymerization. DD established wet-dry cycles as sources of chemical potential for polymerization of nucleotides and wrote the first draft. All authors edited the manuscript.

The authors declare no competing financial interests.

The data generated during or analysed during the current study are available from the corresponding author on reasonable request.

## Materials and Methods

### Atomic force microscopy

We used a Cypher from Asylum (now Oxford instruments) equipped with a standard AC240 silicon tip from Olympus with a spring constant around 2 nN/nm and a resonant frequency around 70 kHz. Images were at least 512*512 pixels and aimed for a resolution of 1 nm per pixel or less. The scanning was performed in ambient conditions at 1 Hz. The height of the rings (thickness) was measured using the section analysis tool in the Igor pro software for the AFM.

### Wet-dry cycles on glass substrates

The initial experiments used acid forms of two mononucleotides, adenylic acid (AMP) and uridylic acid (UMP) in 1:1 mole ratios. We chose mononucleotides capable of base pairing, expecting that they might produce more stable polymers than single nucleotides. The acid forms of the nucleotides lowered the pH to 2.5 as expected for a 10 mM solution of a weak acid. This pH range simulates the acidity of water circulating in hot springs and favors condensation reactions that link mononucleotides by phosphoester bonds. Experiments with individual mononucleotides were performed as controls. In preliminary experiments the reactions were carried out on microscope slides having three wells into which 0.1 mL of 10 mM nucleotide solutions were added. The slides were placed on a laboratory hot plate set to 80 °C. The solution evaporated in a few minutes, then remained in a dry state for 30 minutes during which phosphoester bonds were expected to form by condensation. Because none of the components could be readily oxidized by molecular oxygen, the experiments were carried out in open air.

After 30 minutes, each well was rehydrated with 50 microliters of water and the reactants were stirred for a few seconds with the end of a cleaned stainless steel spatula. At the end of three such wet-dry cycles 100 uL of water was added and transferred from one well to the next to dissolve any products. This was repeated two more times and each 100 uL sample was placed in a 1.5 mL Eppendorf centrifuge tube. Half of the 300 uL total was treated by a standard ethanol procedure used to precipitate oligonucleotides as pellets which were then dissolved in 50 uL of water. Yields varied from one run to the next but typically amounted to tens of micrograms, representing up to 10% yields of polymers based on the monomers present as reactants.

To determine whether rings were present, a 20 uL aliquot of the cycled reaction mixture with products was placed on freshly cleaved mica for 30 seconds to allow polymers to adhere, then flushed with deionized water. The surface was dried under a stream of nitrogen gas, then examined by atomic force microscopy. This method is also used for studying DNA^28^.

### Experiments on mica surfaces

We added 20-40 µl of the mononucleotide solutions to the surface of freshly cleaved mica which was cut to fit the stage of the AFM. The mica with nucleotide solution was then placed on a laboratory hot plate maintained at 80° C. The solution was allowed to dry for 30 minutes after which the mica surface was rehydrated with 20-40 µl ultrapure deionized water (MilliQ). This was repeated 3 times. After the final dry cycle the mica surface was flushed with ample amounts of water (3-5 ml) and dried under a gentle stream of nitrogen gas. The sample was immediately mounted and examined in the AFM.

**Extended data Fig 1.**
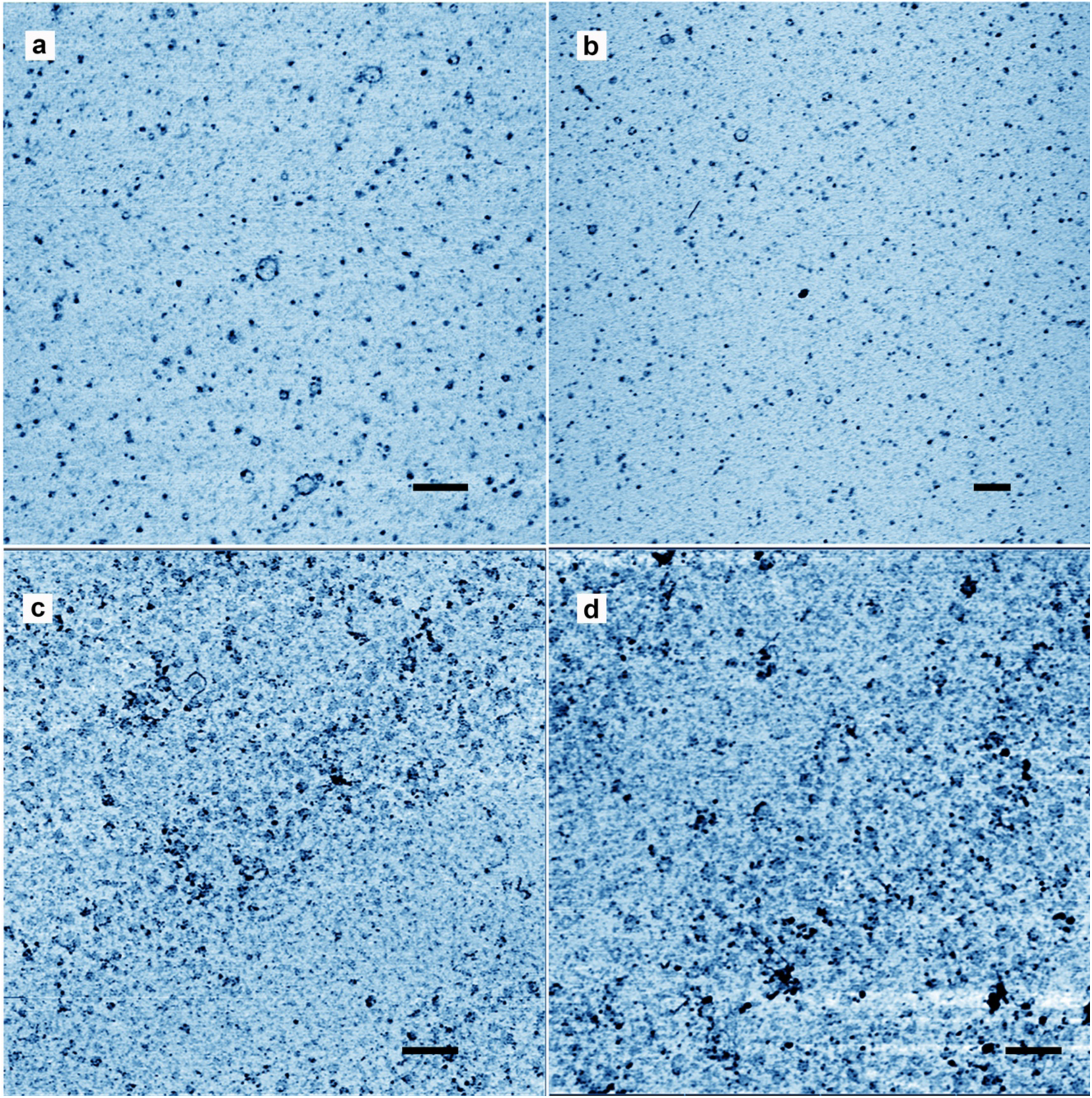
Additional data for AU mixtures. Rings were abundant after wet-dry cycling. The four micrographs show a mica surface which was exposed to a solution of AMP and UMP that had been cycled three times on a glass substrate. The images shown are from two different experiments in two different laboratories. Both display rings, but the amount of material deposited was different, making c) and d) not as clear as a) and b). The scale bars shows 200 nm.

**Extended data Fig 2.**
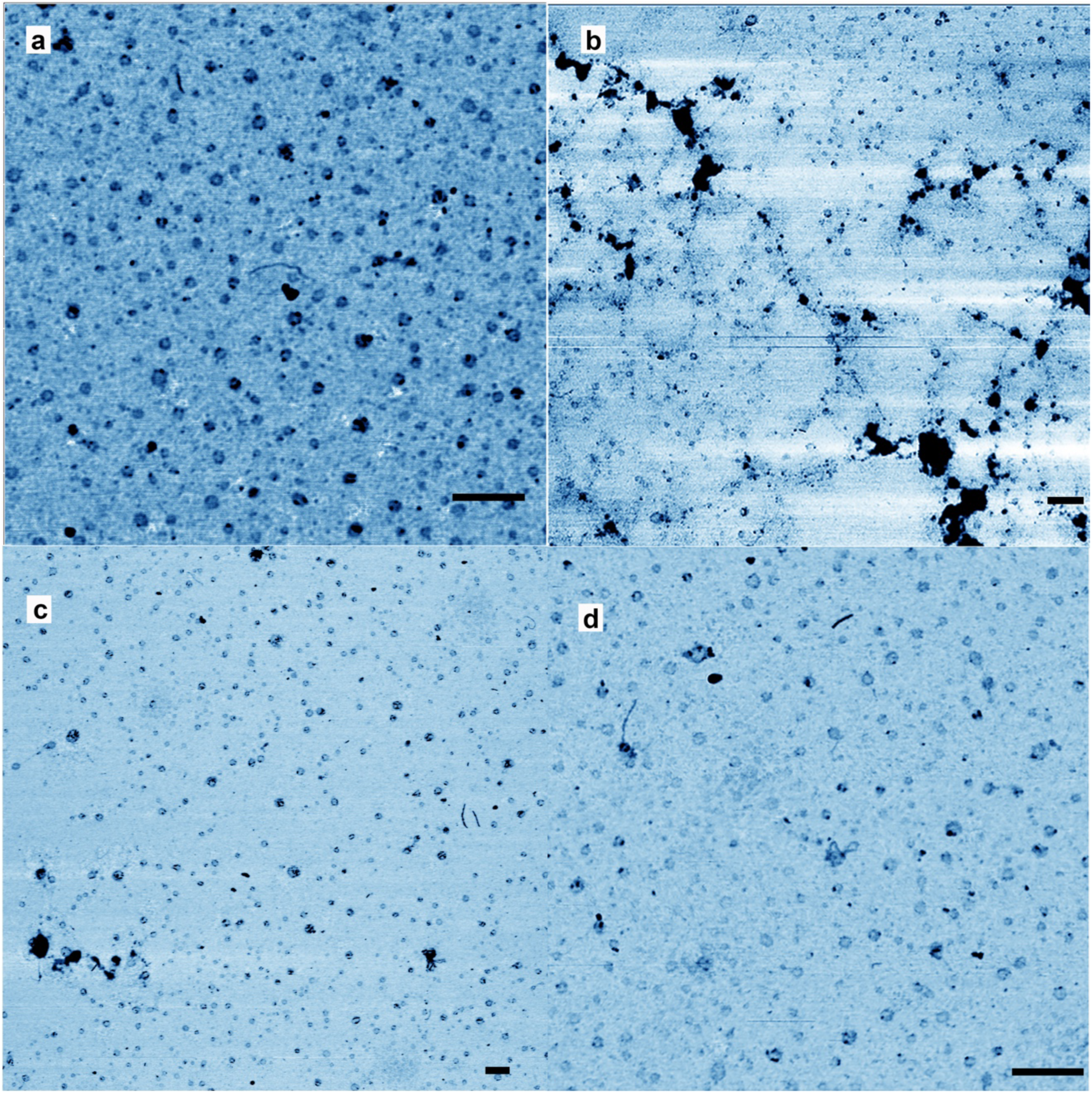
Additional data for GC mixtures. Rings were abundant after wet-dry cycling. The four micrographs show a mica surface on which a mixed solution of GMP and CMP had been cycled three times. The scale bars shows 200 nm.

**Extended data Fig 3:**
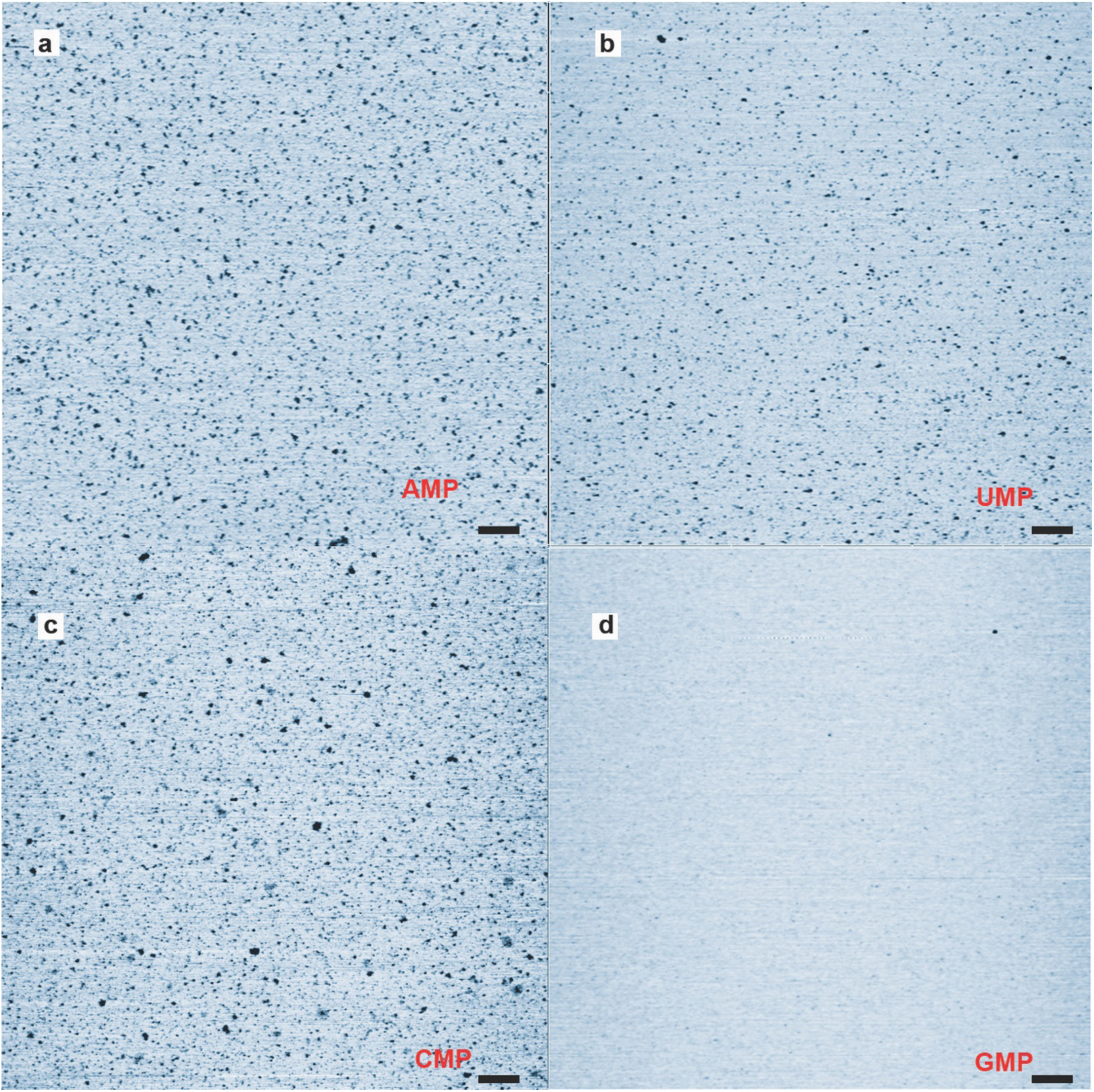
Controls were run with uncycled 10 mM mononucleotides on mica with a) AMP, b) UMP, c) CMP and d) GMP. The protocol was the same as that used for in Extended data Fig. 1. A few microliters of solution were added to a freshly cleaved mica surface, left there for 30 seconds and then rinsed with pure water. Only particles were apparent, showing that the solutions did not contain rings at the outset.. The scale bars shows 200 nm.

**Extended data Fig 4:**
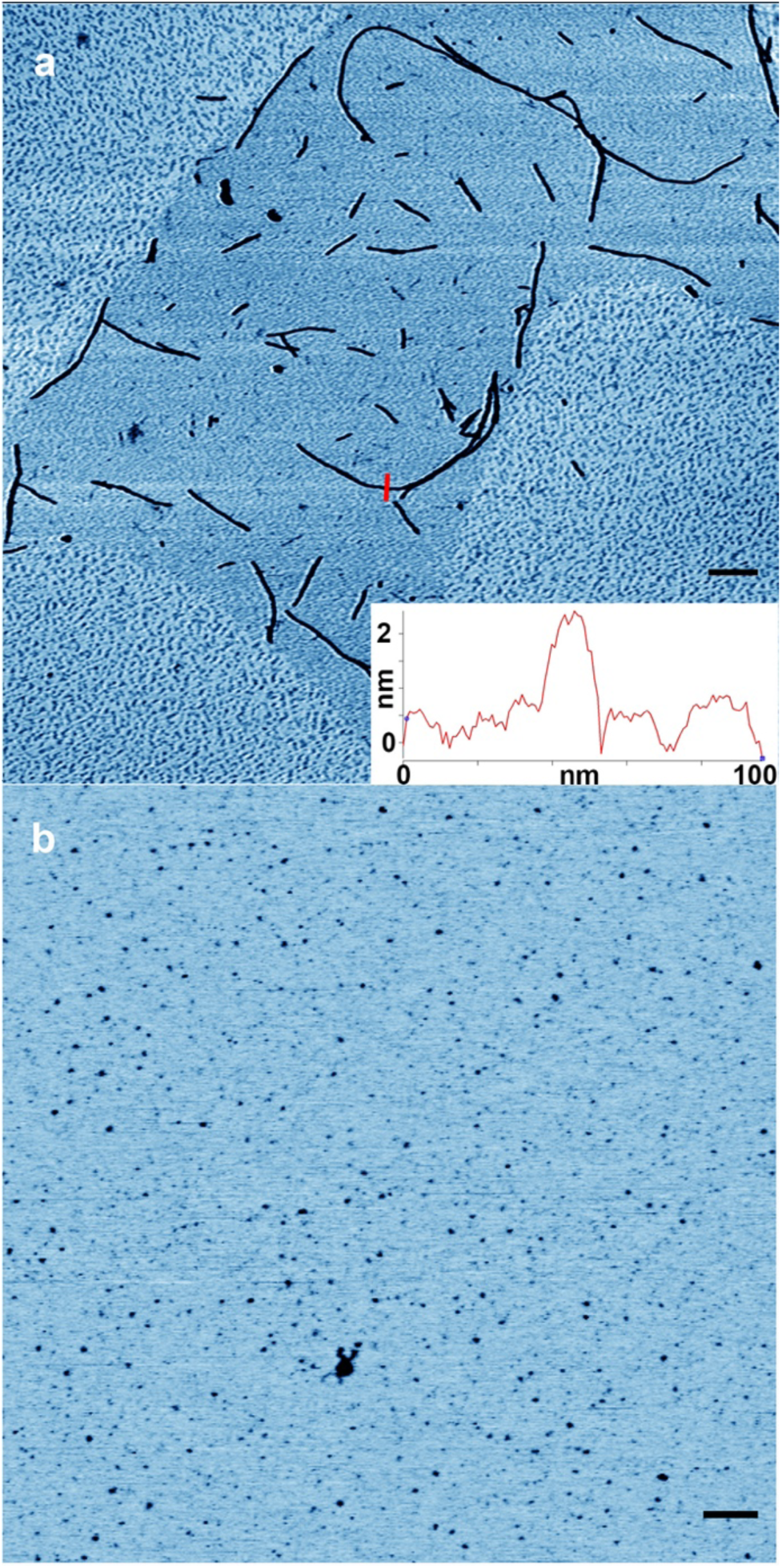
Products from room temperature wet-dry cycling of a) 1:1 GC and b) 1:1 AU mixture solutions. The cross section in a) shows that the GC fibrils are more than 2 nm thick and 14 nm wide, significantly different from the rings observed with wet-dry cycling at 80 °C. The fragmented forms and the mostly linear shapes suggests that the structures are soft crystals rather than polymers. No ring structures were observed with the GC or AU mixtures. The scale bars shows 200 nm.

